# Omicron Subvariants Infection Kinetics and Nirmatrelvir Efficacy in Transgenic K18-hACE2 Mice

**DOI:** 10.1101/2025.07.25.666822

**Authors:** Vijeta Sharma, Enriko Dolgov, Taylor Tillery, Camila MendezRomero, Alberto Rojas-triana, Diana VillalbaGuzman, Kira Goldgirsh, Risha Rasheed, Irene Gonzalez-Jimenez, Nadine Alvarez, Steven Park, Madhuvika Murugan, Andrew M. Nelson, David S. Perlin

**Affiliations:** Center for Discovery and Innovation, Hackensack Meridian Health, 111 Ideation Way. Nutley, New Jersey 07110, United States

**Author notes:** Correspondence should be addressed to David S. Perlin,; and Vijeta Sharma.

**Keywords:** SARS-CoV-2, Omicron Subvariant, Animal Model, Antiviral Agents, Pulmonary immune response, Nirmatrelvir

## Abstract

The persistent evolution of SARS-CoV-2 has led to the emergence of antigenically distinct Omicron subvariants exhibiting increased transmissibility, immune evasion, and altered pathogenicity. Among these, recent subvariants like JN.1, KP.3.1.1, and LB.1 possess unique antigenic and virological features, underscoring the need for continued surveillance and therapeutic evaluation. As vaccines and commercial monoclonal antibodies show reduced effectiveness against these variants, the role of direct-acting antivirals, such as Nirmatrelvir, targeting conserved viral elements like the main protease inhibitor, becomes increasingly crucial. In this study, we investigated the replication kinetics, host immune responses, and therapeutic susceptibility of three recently circulating Omicron subvariants in the K18-hACE2 transgenic mouse model, using the SARS-CoV-2 parent WA1/2020 strain as a reference. Omicron subvariants exhibited a marked temporal shift in viral infection kinetics characterized by an early lung viral titer peak (∼7-8 Log PFU) at 2 days post-infection (dpi), followed by a decline (1–3 Log PFU) by 4 dpi. Pulmonary cytokine and chemokine responses (GM-CSF, TNF-α, IL-1β, IL-6) showed an earlier increase in subvariant-infected mice compared to a gradual response in WA1/2020 infection. Notably, Nirmatrelvir treatment led to significant reduction in lung viral titers in subvariant-infected mice than in WA1/2020, surpassing its efficacy against the parent strain. These findings highlight that Omicron subvariants infection yields a broad dynamic range in viral burden with minimum variability, while retaining a prominent therapeutic response to Nirmatrelvir. This study provides insights to the emerging subvariants pathogenesis and therapeutic responsiveness, reinforcing the importance of continued variant monitoring and the development of effective countermeasures.

**Importance:** The emergence of SARS-CoV-2 Omicron subvariants still poses a significant public health challenge due to their antigenic drift, altered pathogenesis, and immune evasion capabilities. This study comprehensively highlights the distinct replication kinetics, immune responses, and therapeutic susceptibility of the recently circulating Omicron subvariants JN.1, KP.3.1.1, and LB.1 using the K18-hACE2 mouse model. We demonstrate that while these subvariants exhibit altered virological profiles and immune activation patterns compared to the parent WA1/2020 strain, they remain susceptible to Nirmatrelvir which is a direct-acting main protease inhibitor. Notably, Nirmatrelvir demonstrated greater in vivo antiviral efficacy in Omicron subvariant infections than in the ancestral WA1/2020 strain. These findings underscore the enduring therapeutic value of protease-targeting antivirals and emphasize the critical need for ongoing variant-specific preclinical assessments.

## 1. Introduction

Severe acute respiratory syndrome coronavirus 2 (SARS-CoV-2) continues to evolve as one of the most significant public health threats of the 21st century. Since its emergence, it has caused widespread morbidity and mortality, with over 7.78 million deaths globally reported as of December 2024 (WHO report 2024) [1]. A key driver of the virus’s persistence has been its remarkable evolutionary capacity, fueled by the intrinsic infidelity of its RNA-dependent RNA polymerase, leading to the continuous emergence of novel variants with enhanced transmissibility and immune escape properties [2,3].

Among these, the Omicron lineage has proven particularly adept at evading immunity, rapidly diversifying into a multitude of subvariants [4]. Several of these subvariants such as JN.1, KP3.1.1, and LB.1, have risen to dominance between August 2023 and March 2025, displacing earlier strains and posing fresh challenges for both natural and vaccine-induced immunity [5–7]. These subvariants carry extensive spike protein mutations, some of which confer characteristics reminiscent of earlier variants like Delta, including increased fitness and altered tissue tropism [4]. JN.1, a descendant of BA.2.86, is characterized by the L455S mutation in the receptor-binding domain (RBD) of the spike protein, enhancing its binding affinity to the ACE2 receptor and contributing to its rapid global spread [8]. JN.1 contains more than 70 mutations in the spike protein compared to the ancestral Wuhan-Hu-1 strain [9]. As of early 2024, JN.1 accounted for approximately 62% of circulating variants in the United States and was designated a variant of interest by the World Health Organization [10]. In mid-2024, the emergence of new subvariants KP.3 and LB.1, both derived from JN.1, was followed by the evolution of KP.3.1.1 from KP.3 through the accumulation of additional mutations [5,6]. KP.3.1.1 became the predominant variant in the United States, representing over 50% of cases by September 2024. This subvariant harbors mutations such as F456L and Q493E in the RBD, and a deletion at position S31 in the N-terminal domain (NTD), which may alter spike protein conformation and enhance immune escape [11,12]. LB.1, another emerging subvariant, shares similarities with the FLiRT variants and includes a deletion at residue 31 outside the RBD [12]. In August 2024, LB.1 accounted for approximately 14.9% of new cases in the United States. The rapid antigenic shift is thought to be driven by selection pressures from widespread vaccination and repeated natural infections, prompting the virus to find new pathways of immune evasion. While much of the global response has focused on neutralizing antibody-based strategies targeting the spike protein, the pace of antigenic drift has significantly reduced the effectiveness of such countermeasures [12–16].

Importantly, all three subvariants share a conserved P132H mutation in the main protease (Mpro), a mutation located at the interface between domains II and III, a region critical for Mpro structural stability but distal to its catalytic dyad and active site. Biochemical and structural studies have shown that while P132H reduces thermal stability of the enzyme slightly, it does not impair catalytic activity nor compromise binding affinity for selective inhibitors such as Nirmatrelvir [12,17]. Nirmatrelvir, binds to a highly conserved active site unaffected by the P132H mutation, and has emerged as a cornerstone of outpatient COVID-19 treatment due to its broad-spectrum activity and clinical effectiveness [17,18]. Given the declining efficacy of commercial monoclonal antibodies and neutralizing vaccines against these subvariants, small-molecule antivirals directed at conserved viral targets, including Mpro, methyltransferases and RNA polymerase, offer a more resilient therapeutic approach [17–19]. However, with the virus in a state of constant evolution, there remains an urgent need to evaluate the effectiveness of such antivirals against newly emerging subvariants. Understanding the infection dynamics and therapeutic susceptibilities of these subvariants is crucial for informing public health strategies and treatment approaches.

In this study, we investigated the viral replication dynamics, immune response, and antiviral susceptibility of the currently circulating Omicron subvariants JN.1, KP3.1.1, and LB.1 isolated from New Jersey clinical isolates, using the K18-hACE2 transgenic mouse model, a well-established platform for preclinical SARS-CoV-2 pathogenesis and intervention studies [20,21]. Specifically, we investigated whether these subvariants exhibit altered infection kinetics, including early viral peaks and altered host immune response, relative to the parent WA1/2020 strain. Furthermore, we assessed the in vivo efficacy of Nirmatrelvir against these subvariants to test their therapeutic susceptibility and the hypothesis that the K18-hACE2 mouse model provides a robust platform for preclinical antiviral evaluation across emerging variants integrating virological, immunological, and therapeutic assessments. This work provides valuable insights into the ongoing evolution of SARS-CoV-2, the continued utility of the transgenic K18-hACE-2 mouse model for efficacy evaluation and the importance of developing antivirals with novel targets in the face of continued viral diversification.

## 2. Results

### 2.1 Altered lung infection kinetics of Omicron subvariants in K18-hACE2 mice compared to parent WA1/2020

To evaluate the infection dynamics of emerging Omicron subvariants, we intranasally inoculated K18-hACE2 transgenic mice with JN.1, LB.1, KP3.1.1 or the parent WA1/2020 strain at doses shown in **Fig. 1a**, table 1. To assess clinical disease severity, we monitored body weight in K18-hACE2 mice over 4 days post-infection (dpi) with WA1/2020 or Omicron subvariants JN.1, LB.1, and KP.3.1.1. Mice infected with WA1/2020 showed a progressive decline in body weight, culminating in an average loss of ∼10% by 4 dpi. In contrast, mice infected with JN.1 maintained baseline weights throughout the observation period, with no net loss observed. LB.1 and KP.3.1.1 infected mice exhibited modest fluctuations, with average body weights remaining above 95% of baseline on all days with 2-3% net weight loss by day 4. These data indicate that Omicron subvariants induce milder clinical phenotypes compared to the parent WA1/2020 strain (**Fig. 1b**).

**Fig. 1.**
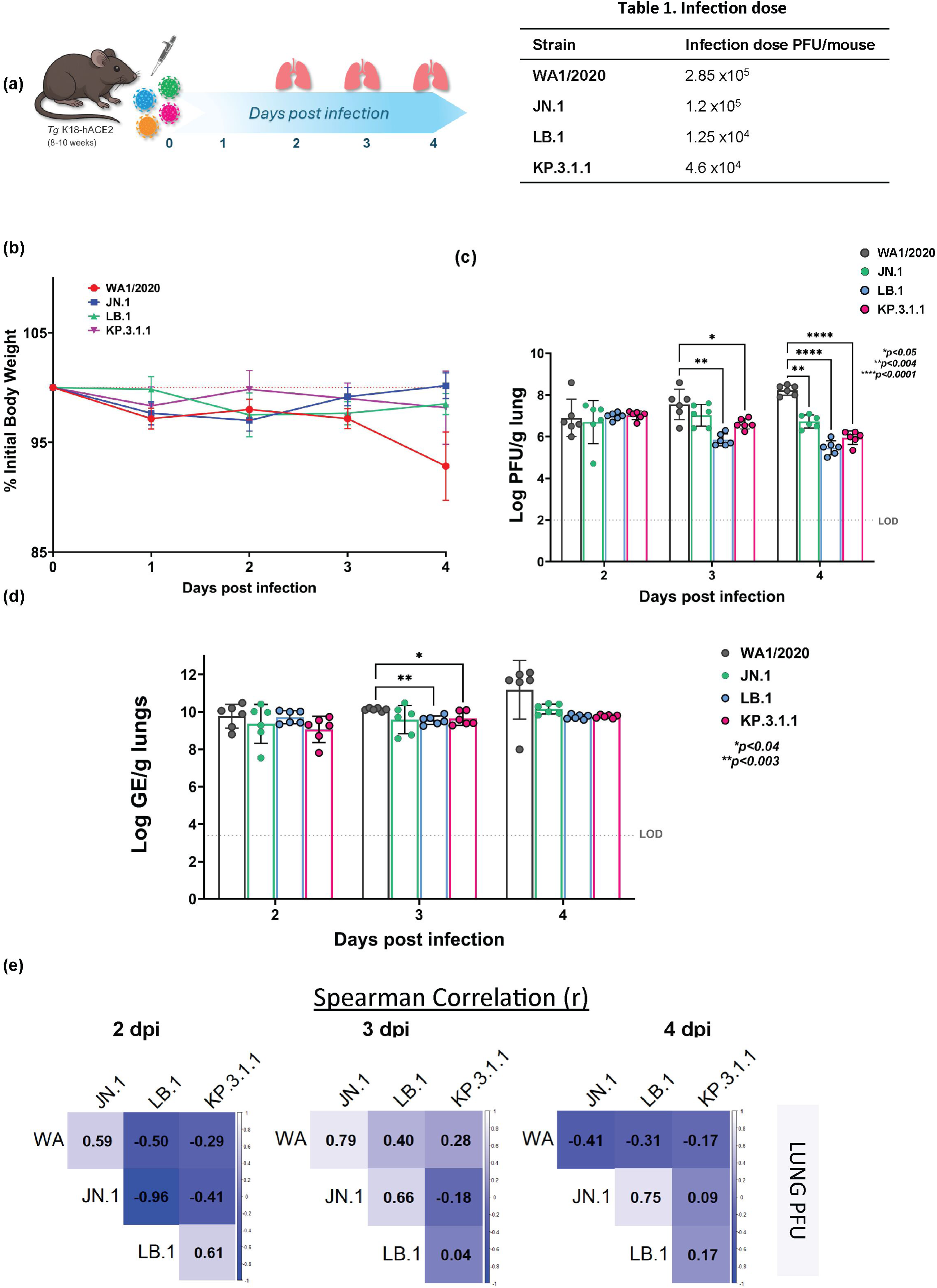
Omicron Subvariants (JN.1, LB.1, KP.3.1.1) infection kinetics in K18-hACE2 mice determined at days 2, 3 and 4 post-infection relative to parent SARS-CoV-2 WA1/2020 strain. (a) Schematic showing the experimental timeline for SARS-CoV-2 infection in K18-hACE2 mice. Male (n=3) and female mice (n=3) K18-hACE2 transgenic mice were infected with parent strain WA1/2020 or Omicron subvariants (JN.1, LB.1, and KP.3.1.1) intranasally at infection doses shown in **table 1** on day 0. Mice were monitored daily. Lungs were collected at necropsy at 2, 3, and 4 days post-infection. (b) Line graphs showing the percent weight loss (% mean ± SEM) in Omicron subvariants JN.1 (blue), LB.1 (green), KP.3.1.1 (purple) and parent WA1 strain (red) infected mice over initial body weight. Bar graphs showing lung viral burdens individual data points in parent WA1/2020 or Omicron subvariants (JN.1, LB.1, KP.3.1.1) infected K18-hACE2 mice at days 2, 3 and 4 post-infection represented as Log_10_ PFU/gram of lung (left) determined by plaque assay (c) and Log_10_ GE/gram of lung (right) determined by qRT-PCR (d). Burden data are presented as mean ± SD with individual data points overlaid. (e) Spearman correlation matrices calculated in R, showing the relationships between lung viral burden (Log PFU/g) in K18-hACE2 mice infected with SARS-CoV-2 WA1/2020 and Omicron subvariants (JN.1, LB.1, KP.3.1.1) at days 2 (left), 3 (middle), and 4 (right) post-infection. Each cell represents the Spearman’s rank correlation coefficient (ρ) between the viral burden of the variant indicated in the row versus the variant indicated in the column. The dashed line indicates the limit of detection (LOD) and p values are mentioned on the graph. Statistical significance was determined by two-way ordinary ANOVA with Sidak’s *post-hoc* multiple comparisons.

To evaluate viral replication kinetics in the lower respiratory tract, lung viral load was assessed as Log PFU/g of tissue by plaque-forming unit (PFU) assay and Log GE (genomic equivalents)/g of tissue by RT-PCR at days 2, 3, and 4 post infection in K18-hACE2 mice (**Fig. 1c, d**). WA1/2020-infected mice showed high lung titers across all days, with mean Log PFU values ranging from ∼7.0 to 8.5, attaining peak titer at 4 dpi (**Fig. 1c**). Mice infected with JN.1 exhibited moderately lower titers at all time points, with mean Log PFU levels 6.7–7 across days 2-4. While stable, the day 4 titers were significantly lower than those observed in WA1/2020, indicating a degree of attenuation in lung viral replication. LB.1-infected mice showed reduction in viral titers. On day 2, mean lung viral titers were Log 7.0 PFU, but decreased gradually to mean Log PFU values of 5.9 and 5.5 by 3 and 4 dpi, respectively. KP.3.1.1 displayed similar kinetics, peaking at ∼7.0 Log PFU at day 2, dropping to 6.6 and 5.9 Log PFU at 3 and 4 dpi, respectively. The lung titers revealed no significant differences between WA1/2020 and any Omicron subvariants at 2 dpi. However, by 3 dpi, LB.1 and KP.3.1.1-infected mice had significantly reduced titers compared to WA1/2020, with mean differences of 1.7 log (p=0.0032) and 0.935 log (p=0.0489), respectively, while JN.1 showed a non-significant trend. At 4 dpi, the differences became more pronounced: WA1/2020 vs. JN.1 (1.5 log, p=0.0012), vs. LB.1 (2.767 log, p<0.0001), and vs. KP.3.1.1 (2.285 log, p<0.0001), indicating robust clearance in Omicron-infected lungs. While WA1-infected mice exhibited peak lung viral titers at 4 dpi, all Omicron subvariants exhibited temporal shift, peaking at 2 dpi in LB.1 and KP.3.1.1 infected mice, and 3 dpi in JN.1 infected mice (**Fig.1c)**. This shift suggests accelerated lung viral clearance in Omicron-infected mice. These data indicate that parent WA1/2020 replicates more efficiently in the lower respiratory tract, followed by JN.1, while LB.1 and KP.3.1.1 exhibit attenuated viral replication in lungs consistent with reduced pathogenicity.

To complement the replicating viral quantification by PFU, lung viral RNA levels were also measured by RT-PCR analysis at days 2, 3, and 4 dpi (**Fig. 1d**). WA1/2020-infected mice exhibited the highest mean RT-PCR log values across all time points, ranging from mean Log GE values of ∼10.0 on day 2 to ∼12.0 by day 4, indicating persistent and escalating viral RNA burden in the lungs. JN.1-infected mice showed slightly lower yet sustained levels of viral RNA, with Log GE means ranging from 9.4 – 10.2 across the three days, suggesting moderately efficient replication compared to WA1/2020. LB.1-infected mice demonstrated a more attenuated viral RNA profile. Mean Log GE values on day 2 were ∼9.7, followed by significantly lower ∼9.6 by day 3 (p=0.0029) than WA1/2020, and remained relatively stable by day 4 (∼9.7), indicating limited viral amplification. Similarly, KP.3.1.1-infected mice exhibited intermediate replication. While early mean RNA levels ranging from Log GE values 9 at 2 dpi, stabilized to 9.6 by 3 dpi which were lower than WA1/2020 (p=0.0378) and remained steady through day 4 to 9.8, consistent with a milder viral replication phenotype.

Notably, while WA1/2020 showed continued increase in viral RNA through 4 dpi, Omicron subvariants (JN.1, LB.1, KP.3.1.1) demonstrated plateauing RT-PCR curves after 2 – 3 dpi, further supporting the trend of more rapid viral clearance observed in PFU assays. These findings underscore that WA1/2020 exhibits the most robust replication kinetics in the lower respiratory tract by 4 dpi, while Omicron subvariants especially LB.1 and KP.3.1.1 are associated with attenuated replication, as evidenced by both PFU and RT-PCR measures.

### 2.2 Divergent temporal patterns of lung viral replication among Omicron subvariants and parent WA1/2020

To investigate the relationships in lung viral replication kinetics among SARS-CoV-2 variants, we performed Spearman correlation analyses using plaque-forming unit (PFU) titers at 2, 3, and 4 dpi in K18-hACE2 mice (**Fig. 1e**). At 2 dpi, WA1/2020 and JN.1 showed a moderate positive correlation (ρ=0.59), suggesting aligned replication trends early in infection. In contrast, WA1/2020 was moderately negatively correlated with LB.1 (ρ=-0.50) and weakly negatively correlated with KP.3.1.1 (ρ=-0.29), indicating that these Omicron subvariants may follow divergent replication trajectories from the parent strain. A striking finding was the very strong negative correlation between JN.1 and LB.1 (ρ=-0.96), indicating inverse viral replication trends. Meanwhile, LB.1 and KP.3.1.1 were moderately positively correlated (ρ=0.61), suggesting partial alignment in viral dynamics among Omicron lineages. By 3 dpi, the relationships shifted substantially. WA1/2020 and JN.1 displayed a strong positive correlation (ρ=0.79), indicating synchronous viral replication during peak infection. WA1/2020 also correlated moderately with LB.1 (ρ=0.40) and weakly with KP.3.1.1 (ρ=0.28). JN.1 and LB.1 maintained a moderate positive correlation (ρ=0.66), in stark contrast to their inverse trend at 2 dpi, suggesting that replication kinetics may converge at this intermediate stage. Relationships between JN.1 and KP.3.1.1 were weakly negative (ρ=-0.18), and LB.1 and KP.3.1.1 were weakly correlated (ρ=0.04). At 4 dpi, WA1/2020 showed negative correlations with all other variants, including JN.1 (ρ=-0.41), LB.1 (ρ=-0.31), and KP.3.1.1 (ρ=-0.17), indicating a divergence in viral persistence, as WA1 continued to replicate while Omicron subvariants declined. In contrast, JN.1 and LB.1 showed a strong positive correlation (ρ=0.75), indicating convergence in viral burden reduction. Similarly, KP.3.1.1 showed positive though weaker correlation with JN.1 (ρ=0.09) and LB.1 (ρ=0.17).

Overall, the correlation analysis revealed dynamic and temporally variable relationships among SARS-CoV-2 variants in lung replication. At early timepoints (2 dpi), inverse replication trends were apparent between JN.1 and LB.1, while WA1 aligned most closely with JN.1. As infection progressed, JN.1 KP.3.1.1 and LB.1 became increasingly synchronized, while WA1/2020 began to diverge, maintaining high titers as Omicron subvariants declined. These results underscore the distinct temporal patterns of viral replication among Omicron subvariants relative to the parent strain, highlighting the importance of temporal resolution in assessing viral pathogenesis. A similar analysis was performed for viral RNA levels (**Fig. S1**).

### 2.3 Broad in vivo efficacy of Nirmatrelvir across WA1/2020 and Omicron subvariants

To evaluate the antiviral activity of Nirmatrelvir treatment against the parent WA1/2020 strain and the Omicron subvariants JN.1, LB.1, and KP.3.1, K18-hACE2 mice were infected and treated with vehicle (10% Tween 80) or 1000 mg/kg of Nirmatrelvir orally twice daily for four days, starting 12 hours post-infection (hpi) (**Fig. 2a**). No significant weight change was observed in Nirmatrelvir treated groups in comparison to vehicle control at all day’s post-administration (**Fig. S2**).

**Fig. 2.**
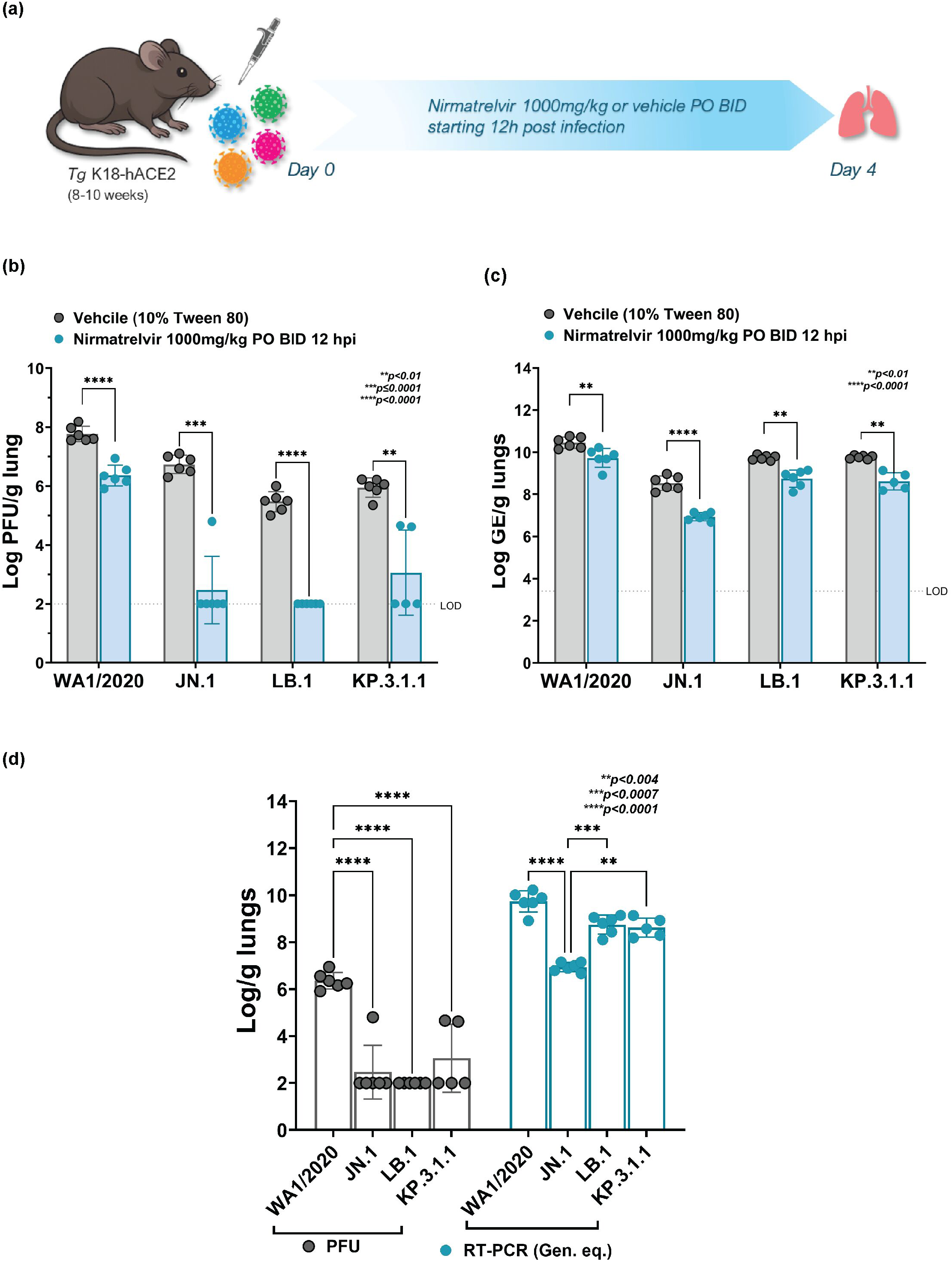
Antiviral response of Nirmatrelvir treatment in Omicron subvariants JN.1, LB.1, and KP.3.1 infected K18-hACE2 mice versus parent SARS-CoV-2 WA1/2020 infected mice. (a) Schematic representation of experimental timeline for Nirmatrelvir treatment in WA1/2020, JN.1, LB.1 or KP.3.1.1 infected K18-hACE2 mice. 8-10 weeks old K18-hACE2 mice (3 male, 3 female, n=6 per group) were intranasally infected with SARS-CoV-2 WA1/2020, JN.1 LB.1 or KP.3.1.1 on day 0 at infection doses mentioned in table 1. Mice were treated orally twice daily (BID) with either Nirmatrelvir (1000 mg/kg) or vehicle 10% Tween 80 (10 mL/kg) starting 12 hpi until day 4 when lung tissue was collected for burden analysis. Lung viral burden in 1000 mg/kg Nirmatrelvir (starting 12 hpi) treated WA1/2020 or Omicron Subvariants (JN.1, LB.1, KP.3.1.1) infected K18-hACE2 mice at days 2, 3 and 4 post-infection represented as Log_10_ PFU/gram of lung (b) determined by plaque assay and Log_10_ GE/gram of lung (c) determined by qRT-PCR. Bars represent mean ± SD, with individual data points shown. The dashed line indicates the limit of detection (LOD). Statistical significance was determined by two-way ANOVA with Tukey’s *post-hoc* multiple comparisons. (d) Comparative analysis of antiviral response of Nirmatrelvir (starting 12 hpi) in WA1/2020 versus Omicron Subvariants (JN.1, LB.1, and KP.3.1.1) infected mice at 4 dpi. Bars represent mean ± SD, with individual data points shown *p* values are mentioned on the graph. Statistical significance was determined by two-way ordinary ANOVA with Sidak’s *post-hoc* multiple comparisons.

Viral load in the lungs was quantified as Log PFU/g and Log GE/g of tissue at the study endpoint (**Fig. 2b, c**). Plaque assay analysis revealed a significant reduction in viral load in the Nirmatrelvir-treated groups compared to the vehicle control across all tested SARS-CoV-2 strains (**Fig. 2b**). Mice infected with the parent WA1/2020 strain showed a significant reduction in mean viral titers from ∼7.8 Log PFU in the vehicle-treated group to 6.53 Log PFU in the Nirmatrelvir - treated group (mean difference = 1.413, p<0.0001).

Mice infected with Omicron subvariant JN.1 demonstrated a substantial response to Nirmatrelvir treatment. The vehicle control group infected with JN.1 showed a mean viral titer of 6.73 Log PFU, which was significantly decreased to 3.22 Log PFU in the Nirmatrelvir-treated group (mean difference = 4.267, p=0.0001). Similarly, Nirmatrelvir treatment showed significant efficacy against the LB.1 and KP.3.1.1 Omicron subvariants. In LB.1 treated mice, the vehicle-treated group showed a mean viral titer of 5.47 Log PFU, which was significantly reduced to 2.50 Log PFU in the Nirmatrelvir-treated group (mean difference = 3.467, p<0.0001). Whereas in KP.3.1.1 infected mice, the vehicle control group showed a mean viral titer of 5.95 Log PFU, which was significantly reduced to 3.83 Log PFU in the Nirmatrelvir - treated group (mean difference = 2.892, p=0.0099).

Consistent with the infectious viral data (PFU), RT-PCR analysis revealed a significant reduction in lung viral RNA levels in the Nirmatrelvir-treated groups compared to the vehicle control across all tested SARS-CoV-2 strains (**Fig. 2c**). In parent WA1/2020 infected mice, the mean viral RNA level in the vehicle-treated group was 10.45 Log GE, which was significantly reduced to 9.85 Log GE in the Nirmatrelvir-treated group (mean difference = 0.72, p=0.0097). The Omicron subvariant JN.1 infected mice demonstrated a marked reduction in lung viral RNA following Nirmatrelvir treatment. The vehicle control group infected with JN.1 exhibited a mean viral RNA level of 8.53 Log GE, which was significantly decreased to 7.13 Log GE in the Nirmatrelvir-treated group (mean difference = 1.59, p<0.0001). Similarly, Nirmatrelvir treatment showed a significant decrease in lung viral RNA levels against the LB.1 and KP.3.1.1 infected mice. In LB.1 infected mice, the vehicle-treated group had a mean viral RNA level of 9.74 Log GE, which was significantly reduced to 8.95 Log GE in the Nirmatrelvir-treated group (mean difference = 0.99, p=0.0012). Finally, in KP.3.1 infected mice, the vehicle control group showed a mean viral RNA level of 9.77 Log GE, while the Nirmatrelvir-treated group showed mean viral RNA level of 8.80 Log GE (mean difference = 1.15, p=0.0027). These findings underscore that Nirmatrelvir retained potent efficacy against Omicron subvariants.

### 2.4 Comparative Nirmatrelvir susceptibility reveal enhanced antiviral response in Omicron subvariants infected mice in comparison to WA1/2020

Nirmatrelvir treatment in K18-hACE2 mice demonstrated significant antiviral activity, as measured by infectious virus as PFU, against both the parent WA1/2020 strain and all Omicron subvariants (JN.1, LB.1, and KP.3.1.1). Nirmatrelvir nearly completely suppressed infectious virus in the Omicron subvariants JN.1 and LB.1, reaching the limit of detection (p<0.0001). KP.3.1.1 also showed a substantial reduction as evident by decrease in PFU (**Fig. 2d**, grey bars) which was significantly lower than parent WA1/2020 (p<0.0001). Nirmatrelvir treatment also significantly reduced viral RNA levels in JN.1 (p<0.0001) treated mice in comparison to WA1/2020 infected mice. However, no significant viral RNA levels reduction was observed in the LB.1 and KP.3.1.1 infected mice following treatment, as compared to the WA1/2020 infected group (**Fig. 2d**, blue bars). Nevertheless, in vitro evaluation of Nirmatrelvir efficacy revealed EC₅₀ values of 0.0158 ± 0.0105 μM for JN.1, 0.060±0.011 μM for LB.1 and 0.090 ± 0.051 μM for KP.3.1.1, compared to 0.0301 ± 0.0114 μM for the WA1/2020 strain (**Fig S3**). Overall, these findings further support the continued susceptibility of Omicron subvariants to Nirmatrelvir treatment, surpassing that was shown in WA1/2020.

### 2.5 Early pulmonary immune response analysis reveal distinct pattern in Omicron subvariants infected mice

To determine the early pulmonary immune activation in response to SARS-CoV-2 infection, cytokine and chemokine levels were quantified in lung homogenates of K18-hACE2 mice at days 2, 3, and 4 post-infection (**Fig. 3a, S4a**). The panel of cytokines and chemokines assessed in this study was chosen based on clinical observations from severe SARS-CoV-2 infected patients, where excessive and dysregulated immune responses— often termed “cytokine storm”, have been implicated in severe disease progression. Previous clinical reports identified several cytokines such as GM-CSF, TNF-α, MCP-1 (CCL2), IL-1β, IL-6, IL-10, CXCL10 (IP-10), and CCL3 (MIP-1α), as key mediators consistently elevated in severe cases, contributing to hyperinflammation, immune-mediated tissue damage, and poor clinical outcomes [23–26]. In this study, these cytokines and chemokines were analyzed in lung homogenates of infected mice to provide a comprehensive understanding of the early immune activation elicited by SARS-CoV-2 infection in the K18-hACE2 mouse model. The levels of the selected cytokines and chemokines were higher from baseline in all the groups suggesting a strong inflammatory response to infection. However, the dynamic range of levels show the heterogeneous nature of the immune response as seen in the patients.

**Fig. 3.**
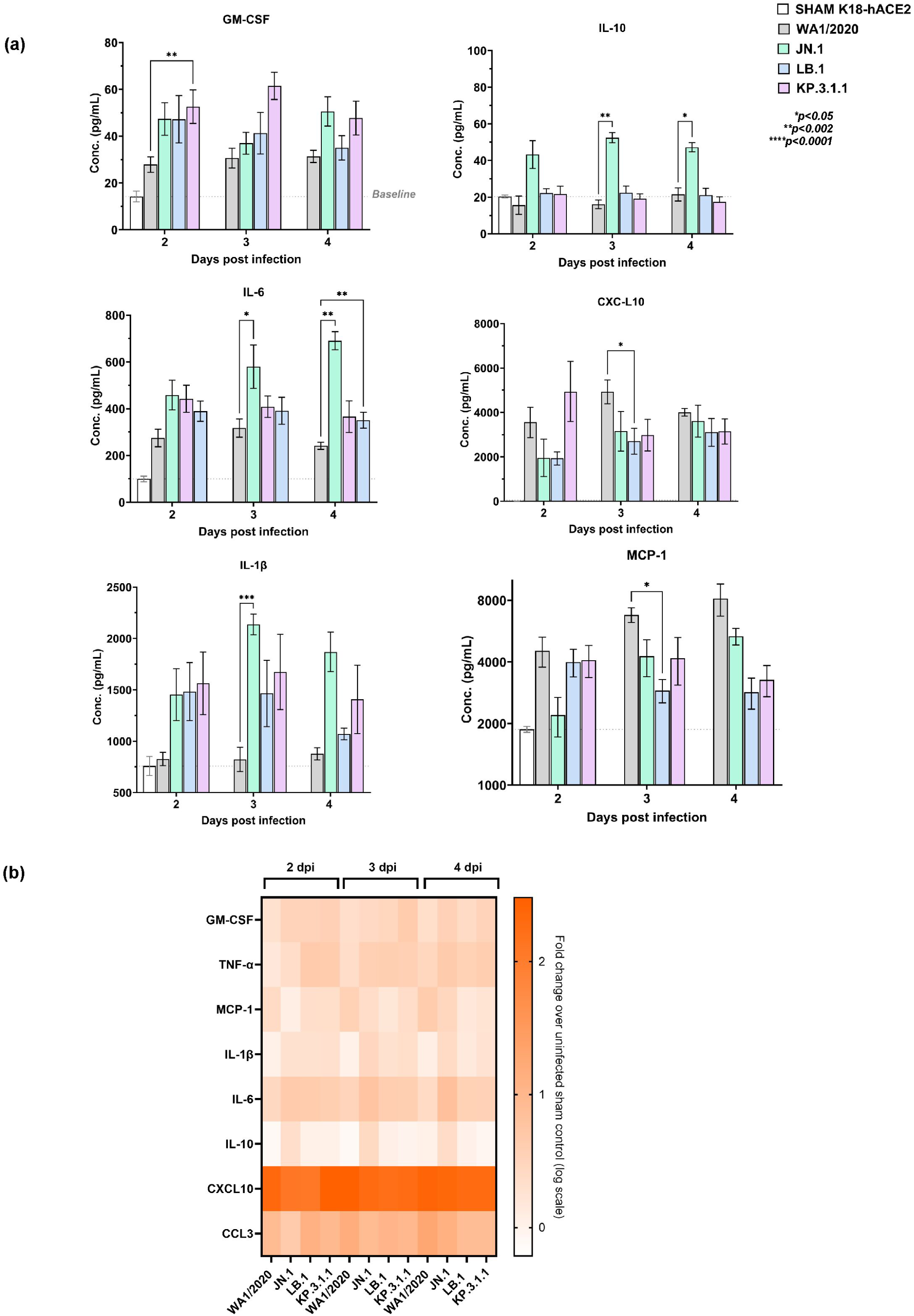
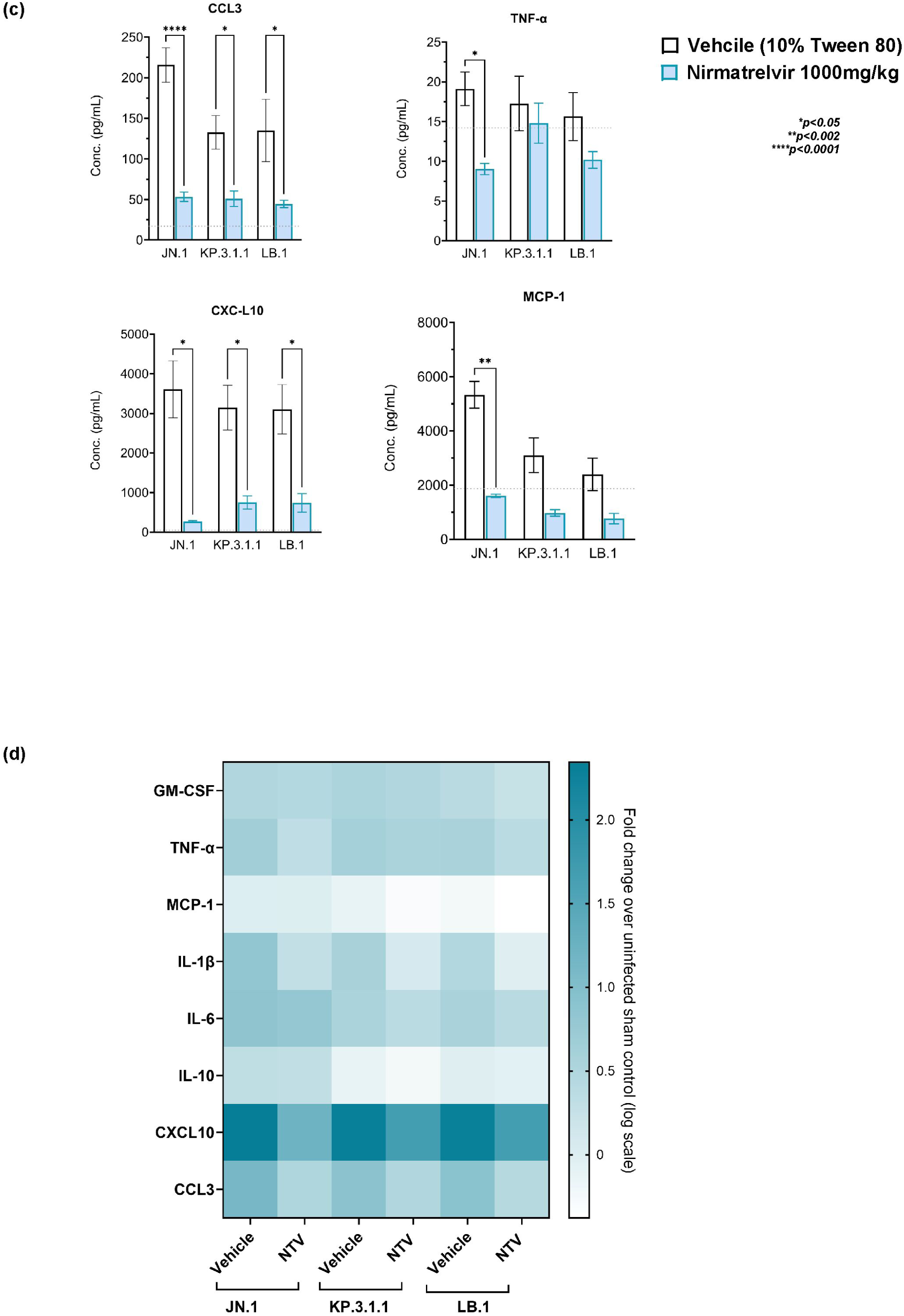
Early pulmonary immune response analysis in Omicron subvariants JN.1, LB.1, and KP.3.1 versus parent WA1/2020 infected K18-hACE2 mice at 2, 3, and 4 days post-infection (dpi). (a) Temporal lung cytokine and chemokine analysis in K18-hACE2 mice following infection. The levels of GM-CSF, IL-10, IL-6, MCP-1, CXCL10, and IL-1β (pg/mL) were measured in lung homogenates at 2, 3, and 4 dpi. Bar graphs represent concentrations as mean± SEM and dashed lines show baseline levels in uninfected sham control. (b) The heatmap illustrates the log_10_ fold change lung cytokines and chemokines levels of K18-hACE2 mice infected with different WA1/2020, JN.1, LB.1, KP.3.1.1 at 2, 3, and 4 dpi, relative to uninfected control mice. The color intensity corresponds to the magnitude of the fold change as indicated by the scale. (c) Effect of Nirmatrelvir treatment (1000 mg/kg) on lung cytokines and chemokine levels in JN.1, LB.1 and KP.3.1.1 infected mice in comparison to the infected vehicle controls. The levels of TNF-⍺, IL-6, MCP-1, CCL3, and CXCL10 were measured in lung homogenates at 4 dpi. Bar graphs represent concentrations (pg/mL) as mean±SEM with dashed lines showing baseline levels in uninfected sham control. (d) The heatmap illustrates the log_10_ fold change in the lung cytokines and chemokines levels of Nirmatrelvir treated K18-hACE2 mice infected with JN.1, LB.1, and KP.3.1.1 versus infected vehicle control. The color intensity corresponds to the magnitude of the fold change as indicated by the scale. *P* values are mentioned on the graph wherever required. Statistical significance was determined by two-way ordinary ANOVA with Sidak’s *post-hoc* multiple comparisons.

At 2 dpi, WA1/2020-infected mice showed moderate GM-CSF induction with a mean of 28 pg/mL, while JN.1 and LB.1 infected mice exhibited a mean value of ∼47 pg/mL, respectively, which was significantly higher (p=0.0087) in KP.3.1.1 infected mice with a mean of 53 pg/mL, reflecting an accelerated early immune activation **(Fig. 3a**). At 3 and 4 dpi, there was a slight increase in the levels with no significant difference in the levels in Omicron infected mice in comparison to WA1/2020.

At 2 dpi WA1/2020 induced mild TNF-α (mean 6.4 pg/mL), which is a proinflammatory cytokine, while JN.1, KP.3.1.1 and LB.1 showed mean of ∼10, 19and 20 pg/mL though not significantly different from the WA1/2020 infection. By 4 dpi, all variants showed a similar trend in TNF-α levels **(Fig. S4a)**.

MCP-1 levels followed a similar trend in all the groups in the range of 2000 - 4000 pg/mL at 2 dpi, which was significantly decreased (p=0.04) in LB.1 infected mice at 3 dpi in comparison to WA1/2020 (**Fig. 3a**). MCP-1 levels showed a gradual increase from mean of 4529 - 8151 pg/mL in WA1/2020 while Omicron infected mice showed a sustained trend of mean 2200 - 5333 pg/mL in JN.1, 3994 - 2800 pg/mL in LB.1 and 4085 - 3272 pg/mL in KP.3.1.1 over 2 - 4 days period. The increasing trend in MCP-1 levels in WA1/2020 infection is indicative of strong monocyte recruitment in lungs, which was mild in Omicron infected mice.

WA1/2020-infected mice had moderate IL-1β levels of mean ∼827 pg/mL, at 2 dpi, while JN.1, KP.3.1.1 and LB.1 infected mice had higher means of 1454, 1564 and 1484 pg/mL, respectively (**Fig. 3a**). At 3 dpi, JN.1 showed significantly higher levels with a mean difference of 1379 pg/mL (p=0.0003) than WA1/2020. There was no significant change in the levels at 4 dpi in all subvariant-infected groups than WA1/2020 showing a more sustained IL-1β response, reflecting a steady immune activation pattern.

IL-6, a central pro-inflammatory cytokine, showed a mean of 275, 459, 443 and 390 pg/mL in WA1/2020, JN.1, LB.1 and KP.3.1.1 infected mice, respectively, at 2 dpi **(Fig. 3a**). By 3 and 4 dpi, IL-6 levels further increased across all groups, indicating persistent immune activation. Nonetheless, JN.1 showed a significant increase to mean 580 pg/mL (p=0.0181) relative to WA1/2020 at 3 dpi. At 4 dpi, KP.3.1.1 and JN.1 showed a significant increase to a mean of 351 pg/mL (p=0.0093) and 691 pg/mL (p=0.0028). Overall, Omicron subvariants infection induced a more pronounced early response than the parent strain.

IL-10 levels showed a minimal increase from baseline IL-10 in all groups at 2, 3 and 4 dpi suggesting a balanced anti-inflammatory response **(Fig. 3a**). This increase was significant in JN.1 infected mice at 3 dpi (p=0.0087) and 4 dpi (p=0.0119) than WA1/2020 which implies JN.1 induces early anti-inflammatory response in lungs.

CXCL10 was markedly elevated in all variants from baseline, which was highest in WA1/2020 infected mice indicating strong interferon-driven responses across 2-4 dpi. However, LB.1 showed a significant decrease (mean difference =2297 pg/mL; p=0.047) than WA1/2020 at 3 dpi. Another chemokine, CCL3, had a similar trend in all the groups through 2 - 4 dpi maintaining strong chemokine production (from baseline) which is indicative of macrophage activation in lungs **(Fig. S4a)**. These results were further analyzed by the heatmap which reveals distinct temporal and variant-specific immune profiles (**Fig. 3b**). The fold change pattern reveals a distinct cytokine and chemokine profile in Omicron subvariant-infected mice where an early immune response was observed, while WA1/2020-infected mice displayed a more gradual and steady immune response.

The observed differences in cytokine and chemokine levels between WA1/2020 and Omicron subvariant-infected mice can be attributed to the distinct replication dynamics and immune response of these viral strains. Elevated levels of GM-CSF, IL-6, IL-1ꞵ, and CXCL10 in Omicron infected mice suggest enhanced early immune activation. Conversely, WA1/2020 maintained a more balanced cytokine and chemokine profile, consistent with its robust lung replication. These findings highlight the distinct immunopathological profiles of Omicron subvariants.

### 2.6 Nirmatrelvir treatment attenuates inflammatory cytokine and chemokine responses in Omicron-infected mice

To assess the immunomodulatory impact of Nirmatrelvir, we quantified cytokines and chemokines levels in lung homogenates of K18-hACE2 mice infected with Omicron subvariants (JN.1, LB.1, KP.3.1.1) and treated with Nirmatrelvir or vehicle. Results revealed distinct suppression of key inflammatory mediators with treatment. At day 4 post-infection, several key differences in lung cytokine and chemokine levels were observed between the vehicle-treated and Nirmatrelvir-treated groups (**Fig. 3c, Fig. S4b**). Nirmatrelvir treatment noticeably suppressed the pulmonary inflammatory response. The levels of GM-CSF, TNF-⍺, IL-6, MCP-1, CCL3, CXCL10, and IL-1ꞵ showed a marked decrease in Nirmatrelvir treated mice. Among these, TNF-α was significantly reduced in JN.1-infected mice by mean ∼10.1 pg/mL following Nirmatrelvir treatment (p=0.011), while KP.3.1.1 and LB.1 showed no significant change (**Fig. 3c**). A mean 3723 pg/mL reduction in MCP-1 levels was observed in JN.1-infected mice (p=0.0017) upon Nirmatrelvir treatment. Though low, changes in MCP-1 levels in LB.1 and KP.3.1.1 were not statistically significant with mean differences of 1632 pg/mL and 2131 pg/mL respectively (**Fig. 3c**). CXCL10 was significantly suppressed in all subvariant-infected groups with Nirmatrelvir treatment which was reduction by mean of 3335 pg/mL in JN.1 (p=0.0169), 2396 pg/mL in KP.3.1.1 (p=0.0207), and 2362 pg/mL in LB.1 (p=0.0326) in comparison to vehicle treated infected controls. Similarly, Nirmatrelvir treatment robustly suppressed CCL3 across all groups, showing the strongest downregulation in JN.1-infected mice with mean difference of 162.3 pg/mL (p<0.0001), which was 81.88 pg/mL (p = 0.0364) in KP.3.1.1 and 90.57 pg/mL (p=0.0133) in LB.1.

While GM-CSF and IL-6 displayed numerically lower levels in Nirmatrelvir treated groups as compared to vehicle treated but the differences were not statistically significant across any subvariant (**Fig. S4b)**. Other cytokines including IL-1β and IL-10 also showed no significant change following treatment, suggesting their expression may be less sensitive to early Nirmatrelvir-mediated modulation (**Fig. S4b)**. The heatmap visualization (**Fig.3d**) further corroborated these findings, illustrating a general downregulation of the fold change in these inflammatory mediators in JN.1, KP.3.1.1 and LB.1 infected mice treated with Nirmatrelvir relative to vehicle-treated infected controls. These findings further support the antiviral potential of Nirmatrelvir and its potential to mitigate the inflammatory response associated with Omicron subvariant infection.

## Discussion

The rapid emergence of SARS-CoV-2 Omicron subvariants has posed ongoing challenges for disease control due to their antigenic drift, altered pathogenesis, and immune evasion capabilities [4–6, 13]. Our study demonstrates that the recent Omicron subvariants JN.1, LB.1, and KP.3.1.1 exhibit distinct infection kinetics and immune responses in the K18-hACE2 mouse model, highlighting variant-specific temporal patterns of viral replication and cytokine activation. Importantly, our results show that these subvariants, despite their altered infection kinetics relative to the WA1/2020 strain, retain robust susceptibility to main protease-targeting antivirals such as Nirmatrelvir [17–22]. A key observation in our study is the accelerated viral kinetics of Omicron subvariants, which displayed peak replication in the lungs at 2 – 3 dpi, followed by a marked decline by 4 dpi. This contrasts with the WA1/2020 strain, which exhibited delayed peak replication and persistent viral RNA accumulation through 4 dpi. Additionally, the Omicron subvariants displayed an enhanced tropism for bronchiolar epithelial cells, as previously demonstrated in our group, in an air–liquid interface (ALI) model using human bronchial airway epithelial cells (hBAECs), where robust infection was observed at 2 and 3 dpi in Omicron infected cells compared to the WA1/2020 strain *ex vivo* [23]. These differences are further corroborated by the variant-specific cytokine and chemokine expression profiles.

Omicron-infected mice mounted an early and transient inflammatory response, specifically characterized by elevated GM-CSF, MCP-1, TNF-α, IL-6, and IL-1β levels [24–27]. Omicron subvariants exhibited an accelerated immune response, characterized by an early peak in cytokine and chemokine levels at 2 - 3 dpi. This rapid immune activation is consistent with the reported upper respiratory tract tropism of Omicron, where robust innate immune sensing occurs early, leading to rapid cytokine production [11]. A similar trend was observed in the ALI model using hBAECs, where a more pronounced immune response was observed in Omicron infected cells compared to the WA1/2020 strain at 2 and 3 dpi [23]. The early peak suggests that Omicron efficiently triggers pattern recognition receptors (PRRs) in airway epithelial cells and alveolar macrophages, leading to a surge in pro-inflammatory mediators [27,28]. However, the faster viral clearance in Omicron-infected mice is accompanied by a decline in immune activation, reflecting the transient nature of this response.

In contrast, WA1/2020-infected mice displayed a gradual increase in cytokine and chemokine levels, peaking around 4 dpi. This delayed response aligns with the stronger lower respiratory tract tropism of WA1/2020, where viral replication in the lungs triggers a slower, more sustained inflammatory response. As WA1/2020 efficiently replicates in lung tissues, immune activation is progressively amplified, leading to peak cytokine levels later in the infection. Additionally, the slower replication dynamics of WA1/2020 may delay antigen presentation and immunity [9,14]. Thus, the temporal differences in immune activation reflect the inherent differences in viral replication kinetics, tissue tropism, and immune activation pathways between the parent strain (WA1/2020) and the rapidly replicating, immune-evasive Omicron subvariants.

The therapeutic component of our study affirms that Nirmatrelvir remains broadly effective against contemporary Omicron subvariants. Treatment with Nirmatrelvir (12 hpi) resulted in significant reductions in both viral titers and RNA levels in the lungs of infected mice, with particularly strong antiviral effects observed in JN.1 and LB.1 infections, reaching near-complete suppression of infectious virus. Interestingly, the antiviral efficacy was more pronounced in subvariant infections than in WA1/2020, potentially reflecting lower baseline replication or potential altered pharmacodynamics in the context of variant tropism [17,18]. Importantly, despite their extensive divergence in spike protein mutations, the Omicron subvariants JN.1, KP.3.1.1, and LB.1 retain the conserved P132H substitution within the main protease, a defining feature of the broader Omicron lineage. This mutation, though changes thermal stability of the enzyme, does not affect the binding activity of selective inhibitors against Mpro like Nirmatrelvir, MK-7845 [18,30,31]. Consistent with these findings, our *in vivo* data show that Nirmatrelvir retains potent antiviral efficacy against all three subvariants.

In addition to reducing viral burden, Nirmatrelvir attenuated the pulmonary cytokines and chemokines response, with notable decreases in pro-inflammatory mediators such as CXCL10, CCL3, and TNF-⍺, MCP-1, supporting its dual role in antiviral suppression and immunomodulation. Critically, our findings validate the continued relevance of the K18-hACE2 mouse model as a preclinical platform for evaluating therapeutic efficacy across emerging SARS-CoV-2 variants. Despite attenuated disease manifestations, Omicron subvariants induced a sufficiently broad dynamic range in viral burden and immune activation, allowing discrimination of therapeutic outcomes. The consistency in infection kinetics and therapeutic response among Omicron subvariants supports the utility of the transgenic mouse model for antiviral efficacy testing.

Collectively, this study advances our understanding of subvariant-specific infection dynamics and therapeutic susceptibility, emphasizing the importance of adaptable preclinical platforms and the continued susceptibility of direct-acting antivirals targeting conserved viral proteins. In the context of rapid viral diversification and declining effectiveness of neutralizing antibody strategies, small-molecule antivirals such as Nirmatrelvir remain critical pillars of SARS-CoV-2 management. Continued genomic surveillance, coupled with timely preclinical evaluation, is essential for informing therapeutic strategies against the evolving viral threats.

## 4. Materials and Methods

### 4.1 Cell and Virus culture

Vero E6 (African green monkey kidney cells) expressing TMPRSS2 (transmembrane serine protease 2) were obtained from XenoTech, Japan (JCRB1819, Lot no. 2222020). The cell monolayers were maintained at 37°C with 5% CO_2_ and 90% relative humidity, in high-glucose Dulbecco’s Modified Eagle Medium (DMEM, ATCC Cat. no. 30-2002TM), supplemented with 10% fetal bovine serum (FBS USDA sourced, Cat. no. FB5002, Thomas Scientific, NJ, US) and 1x antibiotic-antimycotic (AB/AM, Gibco, US; Cat. no. 15240062). The SARS-CoV-2 parent strain (Isolate USA-WA1/2020, Cat. no. NR-52281) was sourced from BEI Resources (NIH supported program managed by ATCC). The SARS-CoV-2 Omicron strains were obtained from BEI Resources and through the Hackensack Meridian Health Bio-Repository (HMH-BioR). All HMH-BioR strains were recovered from nasopharyngeal swabs collected by the New Jersey Department of Health (NJDOH) surveillance program for COVID-19 and were confirmed as SARS-CoV-2 by whole-genome sequencing performed at the New York Genome Center for their unique mutations. Virus SARS-CoV-2 WA1/2020 and Omicron subvariants were propagated in VeroE6/TMPRSS2 cells to prepare stocks. The virus stock titers were determined by plaque assay as plaque forming units per mL of culture as described previously [29]. The stocks used had virus titers of (i) WA1/2020: 5.7 x 10^6^ PFU/mL, (ii) JN.1: 2.4 x 10^6^ PFU/mL, (iii) LB.1: 2.5 x 10^6^ PFU/mL and (iv) KP.3.1.1: 9.2 x 10^5^ PFU/mL.

### 4.2 In vitro antiviral assay

Nirmatrelvir used for *in vitro* and *in vivo* studies was purchased from BOC Biosciences. (CAS. No. 2628280-40-8, Batch no. B22LN0.131). The compound *in vitro* potency was evaluated in VeroE6/TMPRSS2 cells as described previously [30,31]. For the cell-based assay, 10,000 cells per well were seeded in 96-well plates and treated with 3-fold dilutions of Nirmatrelvir 10 mM DMSO stock (36 µM to 0.0018 µM) for 2 h at 37°C and 5% CO2 where media only and uninfected cells served as controls. Following incubation with compound, the cells were infected with each corresponding virus (WA1/2020, JN.1, LB.1 or KP.3.1.1) at 0.5 multiplicity of infection (MOI) and incubated immediately for 3 days under the conditions described above. At 72 h, 100 µL of CellTiter-Glo® 2.0 reagent (Cat. no. G9243) was added to each well for ATP detection by using the TECAN D300e digital dispenser. The half maximal effective concentration (EC_50_) was calculated using the relative luminescence values with respect to uninfected controls.

### 4.3 In vivo studies

All animal procedures described in this study have been approved by the Institutional Animal Care and Use Committee (IACUC) at HMH-Center for Discovery and Innovation. Male and female transgenic B6.Cg-Tg(K18-ACE2)2Prlmn/J mice, 8-10 weeks old were obtained from The Jackson Laboratories, ME, US [17,30]. The animals were housed in individual ventilated caging (IVC) units and handled under sterile conditions for a minimum of 72h for acclimation with food and water *ad libitum* in the Association for Assessment and Accreditation of Laboratory Animal Care International (AAALAC) accredited the Center for Discovery and Innovation (CDI) Research Animal Facility.

Mice (n=6; 3 male and 3 female) from each group were anesthetized by inhalation of vaporized isoflurane and intranasally infected with 50 µL of the virus stock at day 0, to a final infection dose per mouse of (i) WA1/2020: 2.85 x 10^5^ PFU, (ii) JN.1: 1.2 x 10^5^ PFU, (iii) LB.1: 1.25 x 10^5^ PFU and (iv) KP.3.1.1: 4.6 x 10^4^ PFU. Mice were monitored and weights were recorded daily. At necropsy on days 2, 3, 4 post-infection, mice from each group were euthanized by carbon dioxide inhalation and lung, brain and nasal tissues were collected.

For Nirmatrelvir treatment study, mice were administered with Nirmatrelvir at 1000 mg/kg b.w. (body weight) or vehicle (10 mL/kg) via oral gavage twice daily starting 12 hpi until day 3. On day 4 post-infection, mice from each group were euthanized by carbon dioxide inhalation and lungs were collected. The left lung was weighed and homogenized in a GentleMACS M tube containing 2.5 mL of DMEM containing 2% FBS + 1x AB/AM. A clear supernatant was collected from homogenates upon centrifugation at 4000-5000 rpm for 10 min until clear. Cleared supernatants (0.5 mL) were used for PFU assay and remaining 0.25 mL was inactivated by Proteinase K for 45 min at 65⁰C for qRT-PCR.

### 4.4 Viral Load Assessment

#### 4.4.1 Plaque assay

0.1 mL of the clarified homogenates suspended in DMEM were added to a 24-well plate containing 250,000 VeroE6-TMPRSS2 cells per well directly or 10-fold serial dilutions up to 10^-6^ prepared in DMEM. After inoculation, the 24-well plate is gently rocked for 1h at 37⁰C, followed by addition of 0.5 mL of a pre-warmed overlay mixture [2xMEM + 2.5% Cellulose (1:1)]. The plate was incubated for 72 h at 37⁰C. The media was removed, and plates were fixed with 10% neutral buffered formalin and stained with 0.5% crystal violet and washed with water and enumerated.

#### 4.4.2 RNA extraction and RT-qPCR

Total RNA isolation of Proteinase K and heat inactivated lung homogenate supernatants was performed using the Qiagen Qiacube HT automated mid- to high-throughput nucleic acid purification instrument with the QIAamp 96 Virus QIAcube HT Kit. qRT-PCR was performed on the samples using the E gene (from Charité/Berlin (WHO) protocol primer and probe panel) and RNase P gene (CDC kit) [32]. The primer and probe sequences used are: E gene forward 5’ ACAGGTACGTTAATAGTTAATAGCGT 3’; E gene reverse primer 5’ ATATTGCAGCAGTACGCACACA 3’; E gene probe 5’ FAM-ACACTAGCCATCCTTACTGCGCTTCG-BHQ-1 3’; RNAse P Forward 5’ AGA TTT GGA CCT GCG AGC G 3’; RNAse P Reverse 5’GAG CGG CTG TCT CCA CAA G 3’ and RNAse P Probe 5’ RNAse P Probe 3’.

### 4.5 Chemokine and cytokine analysis

Chemokine and cytokine levels of GM-CSF, TNF-α, MCP-1 (CCL2), IL-1β, IL-6, IL-10, CXCL10, and CCL3 were analyzed in collected lung tissue homogenates using the magnetic bead-based Mouse Luminex® Discovery Multiplex Assay (Cat. no. LXSAMSM, RnD Systems, Biotechne, MN, US). Homogenates were mixed with cell lysis buffer 2 (RnD systems, MN, US, Cat. no. 895347) in 1:1 ratio for 30 min and samples were centrifuged at 4,500 g for 10 min at 4°C. The assay was performed using manufacturer’s instructions and read on a Luminex 200 xMAP platform (Austin, TX, US). Cytokine/chemokine concentrations were calculated as picograms per milligram (pg/mL) of tissue using Belysa® immunoassay curve fitting software.

### 4.6 Statistical analysis

The normalized data obtained from the in vitro assays were plotted as effective response *vs.* concentration. The EC_50_ values were determined using a non-linear regression (curve fit), applying the equation for inhibitor *vs.* response variable slope (four parameters) and 95% confidence interval. One way or two-way Analysis of variance (ANOVA) with Tukey’s or Sidak’s or or Dunnet’s, or Bonferroni *post-hoc* multiple comparisons were performed to compare the means of weight change, viral burden and cytokine and chemokine levels among different experimental groups. Statistical analysis was performed using GraphPad Prism v10.4.1 software (GraphPad, San Diego, CA). Spearman’s correlation and normality testing was conducted using R (version 4.4.3) with the corrplot and MVN packages. *P* values of <0.05 were considered statistically significant.

## Author Contributions

Conceptualization, Methodology, Investigation, Validation, Formal Analysis, Graphics, Data Curation: V.S.; *In vivo* experiments: E.D., T.T., D.V., C.M., A.R and VS; *Ex vivo* experiments: V.S. and K.G; Viral stocks preparation and characterization; in vitro screening assays and data acquisition: I.G., R.R., and N.A.; Writing – Original Draft Preparation: V.S. Review and editing: V.S., M.M., N.A., and D.S.P; Supervision: A.V.M. and D.S.P.; Project Administration: M.M., S.P., A.V.M.; Funding Acquisition: D.S.P.

## Funding

This research was funded by NIH/NIAID, grant number 1U19AI171401, granted to David S. Perlin and Charles M. Rice (Metropolitan Antiviral Drug Accelerator, MAVDA).

## Institutional Review Board Statement

All animal studies were ethically reviewed and carried out in accordance with European Directive 2010/63/EU and the GSK Policy on the Care, Welfare and Treatment of Animals. All animal experiments were approved by the Institutional Animal Care and Use Committee (IACUC) at Hackensack Meridian Health, according to guidelines outlined by the Association for the Assessment and Accreditation of Laboratory Animal Care and the U.S. Department of Agriculture.

## Informed Consent Statement

HMH IRB. No. Pro2018-1022, for collection of viral strains.

## Acknowledgments

We would like to thank everyone who helped with this research. We would like to acknowledge the Research Animal Facility (RAF) at Hackensack Meridian Health Center for Discovery and Innovation (HMH CDI) for their support; Sean Fitzgerald and Fatima Ijaz from the Institutional Biological Safety Committee for their assistance; Hackensack Meridian Health Biorepository (HMH BioR) and the Kreiswirth Lab, CDI for generously providing the NJ virus clinical isolates. We acknowledge Wasifa Hafiz Shah for cell line maintenance, and Padmaja Paderu for her effective management of logistics and resources. SARS CoV-2 USA WA1/2020 strain was obtained through BEI Resources, NIAID, NIH.

## Conflicts of Interest

The authors declare no conflicts of interest.

